# Garcinone C suppressed the proliferation of gastric cancer cells through regulating Hedgehog signaling

**DOI:** 10.1101/2025.05.26.655423

**Authors:** Yimeng Zhou, Jin Tae Kim, Jung Won Kwon, Ga Yeon Lee, Hui Mang Son, Kang Hyuk Lee, Shuai Qiu, Hong Jin Lee

## Abstract

Garcinone C, a xanthone derived from *Garcinia mangostana L*, possesses antioxidant and anti-cancer effects. However, its role in gastric cancer remains unexplored. This study aimed to investigate the effects of garcinone C on gastric cancer cell proliferation and its underlying mechanism. We found that garcinone C suppressed gastric cancer cell growth by inducing G0/G1 arrest and apoptosis in a dose-dependent manner. Furthermore, garcinone C downregulated G0/G1 phase markers Cyclin D1 and p21, as well as apoptosis markers Bax, Bcl-2, cleaved-PARP, and c-caspase 3. Following the previous evidence demonstrated that aberrant Hedgehog (Hh) signaling is implicated in gastric cancer development, we confirmed that inhibiting Gli1/2 reduced the growth of AGS and MKN74 cells. Furthermore, garcinone C exerted similar effects to Gant61, inhibiting Hh signaling by reducing Gli1/2 levels and blocking their nuclear translocation. These findings suggest that garcinone C inhibits gastric cancer proliferation via the Hh signaling, indicating its potential as a therapeutic agent for treating gastric cancer.

## 1. Introduction

Gastric cancer remains a major cause of cancer-related death globally^1^. Although its incidence has been gradually declining in developed countries such as the United States and Australia, it remains high in East Asian and Eastern European countries^2^. Unfortunately, most gastric cancer patients are diagnosed at an advanced stage, leading to poor survival rate^3^. In gastric carcinogenesis, the consumption of smoked, pickled, and salt-preserved foods, as well as foods containing nitrites is linked to a higher risk of gastric cancer, whereas a diet rich in fresh fruits and vegetables can protect against gastric cancer^4^. Numerous signaling pathways, including Hedgehog signaling, Wnt/β-catenin signaling, Notch signaling, EGFR/HER2, and PI3K signaling, as well as the Hippo pathway, play pivotal roles in the tumorigenesis, progression, metastasis, and therapeutic responses of gastric cancer^5,6^. Previous studies have demonstrated that phytochemicals have the potential to prevent gastric cancer. For example, the flavonoid astragalin has been reported to induce apoptosis and inhibit gastric cancer migration and invasion through the PI3K/AKT pathway *in vivo* and *in vitro*, while curcumin, the main component of turmeric, suppresses gastric adenocarcinoma cell migration and invasion via blockade of the Gli1-β-Catenin axis^7,8^. In addition, sulforaphane, extracted from broccoli sprouts, has been proved to suppress the proliferation of gastric cancer by inducing S-phase arrest and apoptosis via p53-dependent manner^9^. Therefore, targeting gastric cancer with natural compounds may be an effective strategy for its prevention.

Xanthones, abundant in mangosteen fruit (*Garcinia mangostana L*.), have demonstrated chemopreventive and therapeutic potential in inhibiting carcinogenesis, including cancer proliferation, invasion, and metastasis^10,11^. For instance, garcinone E has been shown to inhibit migration and invasion in ovarian cancer cells by inducing apoptosis^12^, while α-mangostin has been proven to increase the chemosensitivity of gastric cancer cells by facilitating autophagy^13^. Garcinone C has been reported to inhibit the growth of nasopharyngeal cancer cells through regulating the expression of ATR, Stat3, and 4E-BP1^14^. Besides, garcinone C significantly reduced the cell viability of 22RV1 prostate cancer cells and MDA-MB-231 breast cancer cells^15^. Previous studies also demonstrated that garcinone C can suppress colon tumorigenesis by inducing G0/G1 cell cycle arrest^16^. However, the effect of garcinone C on gastric cancer remains insufficiently explored. These findings underscore the potential of garcinone C as a candidate for further investigation in gastric cancer treatment.

Hh signaling plays a critical role in embryonic development and tissue maintenance^17,18^. When ligands Sonic Hh (Shh), Indian Hh (Ihh), or Desert Hh (Dhh) bind to the transmembrane receptor Patched (Ptch1), Hh signaling will be activated. This binding triggers Ptch1 degradation, releasing Smoothened (Smo), which then translocates to the cilia. As a result, Glioma-Associated Oncogene Family Zinc Finger (Gli) 2 and Gli1 translocate into the nucleus, where Gli1 acts as a key transcription factor enhancing Hh pathway activity^18^. Recently, increasing evidence indicates that dysregulation of Hh signaling contributes to the development of cancer, including gastric cancer^19–21^. Previous studies have found a positive correlation between Gli1 and the epithelial-to-mesenchymal transition (EMT) process in gastric cancer patient specimens^19^. High levels of Shh and Ptch1 are associated with poor prognosis and liver metastasis in gastric cancer patients^22^. Our previous study also suggests the possible binding to Gli1 with garcinone C, leading to Gli1 degradation^21^. Therefore, we hypothesized that garcinone C may inhibit gastric cancer growth by targeting Hh signaling and elucidated the underlying mechanism.

## 2. Materials and methods

### 2.1. Reagents and cell culture

Garcinone C (purity > 98%) was purchased from Chengdu alpha company (Chengdu, China). Gant61 and vismodegib were provided by Millipore (Catalog. No 373401, Darmstadt, Germany) and LC Laboratories (Catalog. No V-4050, Woburn, MA), respectively. SNU-1 (KCLB 00001), AGS (KCLB 21739), NCI-N87 (KCLB 60113), MKN74 (KCLB 80104), MKN45 (KCLB 80103), and MKN1 (KCLB 80101) gastric cancer cells were obtained from Korean cell line bank and cultured with RPMI-1640 supplemented with 10% Fetal Bovine Serum (FBS) and 1% Penicillin-Streptomycin (P/S). All cell lines were incubated at 37 ℃ in a 5% CO_2_ humidified incubator.

### 2.2. 3-(4, 5-dimethylthiazol-2-yl)-2, 5-diphenyltetrazolium bromide (MTT) assay

For cell viability assay, the cells were trypsinized and seeded into 96-well plates at a density of 3000 cells/well in 200 μl culture media. The plates were incubated in a humidified incubator at 37°C. After 24 h, cells were treated with different concentrations of garcinone C (0, 1, 5, 10, 30 μM), Gant61 (0, 1, 5, 10, 30 μM), or vismodegib (0, 1, 5, 10, 30 μM) for 72 h. The number of viable cells was determined by MTT (Catalog. No 475989, Sigma-Aldrich, MO, USA) assay. Cell viability was calculated by comparing to the control.

### 2.3. Colony formation assay

Before treatment, AGS and MKN74 cells (1000 cells/well) were seeded in 12-well plate for 24 h. Then cells were treated with garcinone C, Gant61, or vismodegib for 2 weeks. The media was changed every 3 days during this process. Then, the cells were washed with Phosphate-buffered saline (PBS) twice and fixed with 10% formalin for 10 min, and stained with 0.5% crystal violet for 5 min at room temperature. Finally, the visible colonies were counted. Colony formation rate was calculated by comparing to the control.

### 2.4. Cell cycle analysis

AGS and MKN74 cells (8 × 10^5^ cells/well) were seeded in 6 cm dish for 24 h and treated with garcinone C (0, 0.3, 1, 3 μM). After 24 h treatment, cells were fixed with ice cold 70% ethanol. After fixation, the cells were centrifuged, the supernatant was removed, and they were washed with PBS. Subsequently, cells were resuspended in PBS and treated with 100 μg/mL RNase (Catalog. No 70856, Sigma-Aldrich) and stained by PI (propidium iodide, 1 mg/mL, Catalog. No P4170, Sigma-Aldrich) for 30 min at room temperature (RT) in dark condition. Finally, the samples were then analyzed by flow cytometry using a MuseTM Cell Analyzer (Model 0500-3115 Merck Millipore, Darmstadt, Germany). The quantification of cells in G0/G1, S, and G2/M stages of cell cycle was determined using its embedded fully optimized software modules for cell cycle analysis.

### 2.5. Annexin V-FITC/propidium iodide (PI) apoptosis detection assay

AGS and MKN74 cells (8 × 10^5^ cells/well) were seeded in 6 cm dishes and incubated for 24 h. Staining was carried out by using ApoScan Annexin V-FITC Apoptosis Detection kit (Catalog. No LS-02-100, Biobud, Chungbuk, South Kore). After being treated with garcinone C for 24 h, cells, along with the media, were collected and washed twice with cold PBS. The cell pellet was then resuspended in 100 μL of 1× binding buffer. Double staining was performed by adding 5 µL of Annexin V-FITC and 5 µL of PI to the cell suspension, followed by incubation on ice in the dark. After incubation, 400 μL of binding buffer was added to the suspension to reach a final volume of 500 μL. The samples were analyzed by flow cytometry using a BD FACSAria II (BD Biosciences, CA, USA).

### 2.6. Western blot analysis

Protein samples were immediately transferred to Polyvinylidene fluoride or polyvinylidene difluoride (PVDF) membrane (Catalog. No IPVH00010, Millipore, Darmstadt, Germany) from the sodium dodecyl sulfate polyacrylamide gel electrophoresis (SDS-PAGE) gel. After the transfer, membranes were blocked with skim milk for 1 h at RT and incubated with the primary antibody with gentle agitation overnight at 4 ℃. The membranes were then washed with 1 × Tris-buffered saline with Tween 20 (TBST) 3 times before being incubated with the appropriate secondary antibody with gentle agitation for 2 h at room temperature. The primary antibodies for Cyclin D1, p21, Smo, Suppressor of Fused Homolog (Sufu, Catalog. No 2025S), cleaved caspase 3 (Catalog. No 9661S), and secondary antibodies were purchased from Cell signaling Technology (Danvers, MA, USA). Antibodies for Cyclin-dependent kinase 2 (CDK2, Catalog. No 12790S), Gli1 (Catalog. No sc-515781), Gli2 (Catalog. No sc-271786), BCL2 Associated X (Bax, Catalog. No sc-7480), B-cell lymphoma 2 protein (Bcl-2, Catalog. No sc-7382), cleaved-Poly (ADP-ribose) polymerase (PARP, Catalog. No 9542L), and β-Actin (Catalog. No sc-47778) were purchased from Santa Cruz Biotechnology (Santa Cruz, CA, USA). The membranes were then washed again with 1× TBST 3 times before being developed with Horseradish peroxidase (HRP) substrate (Catalog. No 1721064, Biorad, CA, USA) according to the guideline and the resulting chemiluminescence visualized.

### 2.7. siRNA transfection

Cells were seeded in 6 cm dish (8 × 10^5^ cells/dish) for 24 h and then starved overnight. For knockdown of Gli1 or Gli2, cells were transfected with 20 nM Gli1 or Gli2 siRNA for 24 h. The transfection process was conducted with Fugene HD transfection reagent (Catalog. No E2311, Promega). The transfection efficiency was validated by Western blot.

### 2.8. Confocal analysis

AGS and MKN74 cells were seeded in a 3.5 cm glass bottom dish (2× 10^5^/dish) for 24h. After starvation overnight, cells were treated with garcinone C, Gant61 and vismodegib for 24 h. Next, cells were fixed with 4% paraformaldehyde (PFA) for 30 min at RT after washing with PBS once. Then the cells were blocked with 5% BSA for 1h at RT and then incubated with the primary antibody overnight after washing with PBS twice. Then samples were incubated with the secondary antibody with fluorescent dye for 1 h under dark environment after washing with PBS 3 times. And then the cells were stained with 4’,6-diamidino-2-phenylindole (DAPI) mounting solution (Catalog. No H1200-10, Vector Lab, USA) after removing the secondary antibody and washing with PBS once. Finally, the pictures were taken by confocal microscope (LSM800Airy, Zeiss, Germany) at 400 × magnification.

### 2.9. Isolation of cytosolic and nuclear fractions

To isolate cytosolic and nuclear proteins, the NE-PER™ Nuclear and Cytoplasmic Extraction Reagents Kit (Catalog. No 78833, Thermo Fisher Scientific) was used following the manufacturer’s protocol. Briefly, the cytoplasmic fraction was separated by adding CER1 solution, followed by the addition of CER2 solution for 1 minute. The remaining pellet was then resuspended in NER solution to extract the nuclear fraction. *β*-Actin and Lamin B were used as internal controls for the cytoplasmic and nuclear fractions, respectively.

### 2.10. Statistical analysis

All the data are demonstrated with mean ± standard deviation (SD). Differences between groups were conducted with one-way analysis of variance (ANOVA) post hoc Duncan’s test. p-value less than 0.05 was regarded as significant difference.

## 3. Results and discussion

### 3.1. Garcinone C inhibited the proliferation of gastric cancer cells

Evaluation of six different gastric cancer cell lines revealed that AGS had the highest expression of Gli1, the key positive mediator of Hh signaling, whereas MKN74 had lowest expression of Sufu and Gli3R, two negative mediators of Hh signaling^18^ (Fig. 1A). Consequently, AGS and MKN74 were categorized as Hh signaling-high cell lines, whereas SNU-1 and MKN-1 were Hh signaling-low. Garcinone C (5 μM) strongly suppressed proliferation in Hh-high lines, reducing viability by 88.6 ± 1.7% in AGS and 71.6 ± 3.8% in MKN74, compared to 32.8 ± 2.8% in SNU-1 cells and 34.7 ± 6.4% in MNK1 cells (Fig. 1C). Therefore, further experiments were conducted in AGS and MKN74 cell lines. As shown in Fig. 1D, garcinone C dramatically inhibited the cell viability of AGS and MKN74 cells dose-dependently after 72 h treatment. There were 80.8 ± 0.1 % AGS cells and 70.9 ± 2.6 % MKN74 cells being killed after garcinone C treatment at 5 μM for 72 h. Thus, 0.3, 1 and 3 μM garcinone C were selected for further study. Colony formation results indicated that more than 90% and 50% of colonies were decreased by 1 μM garcinone C in AGS and MKN74 cells, respectively, compared with control group (Fig.1E). These results suggested that garcinone C has a remarkable inhibitory effect on the proliferation of AGS and MKN74 gastric cancer cells.

**Figure 1.**
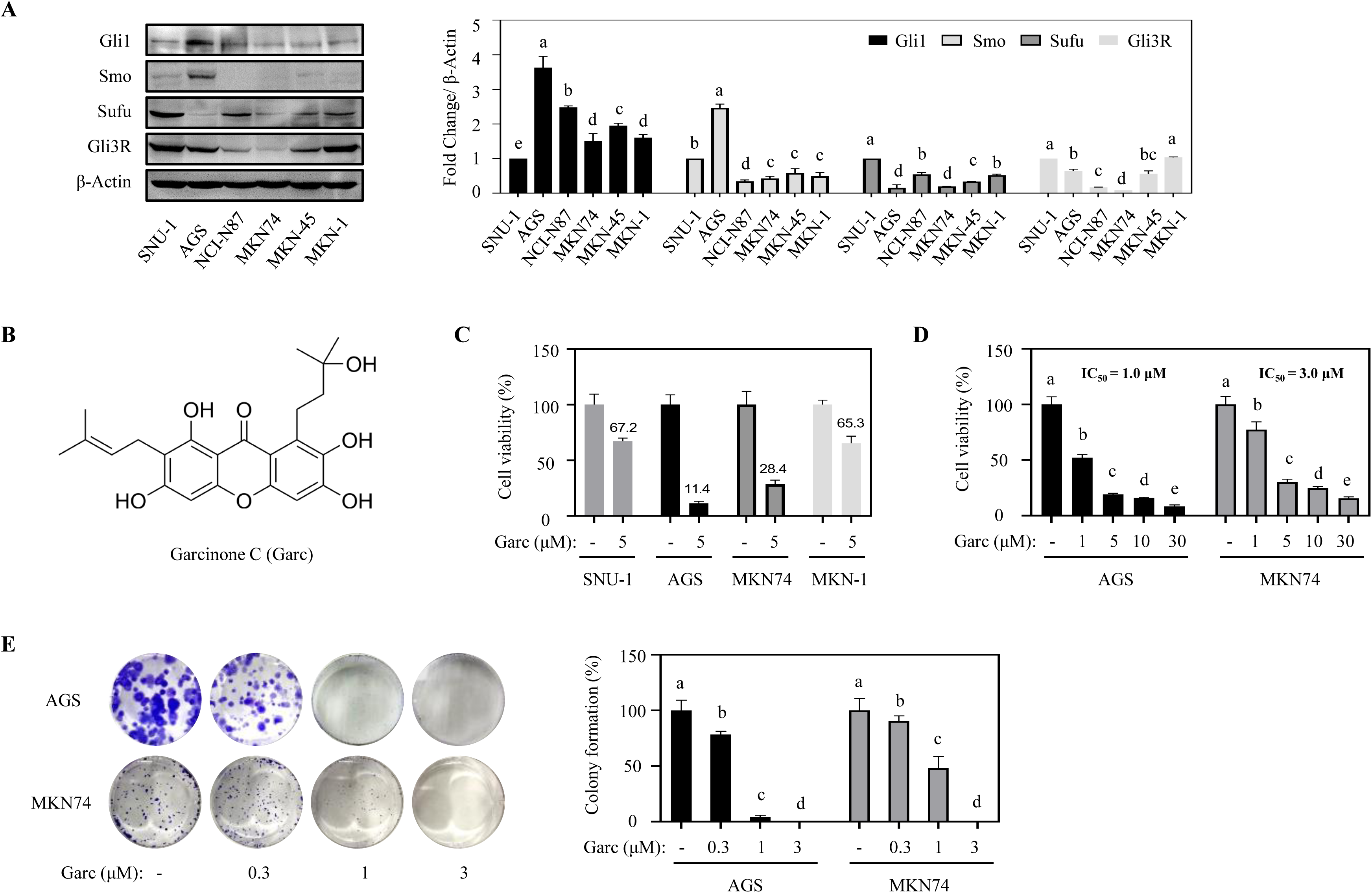
Garcinone C suppressed the proliferation of gastric cancer cells. (A) SNU-1, AGS, NCL-N87, MKN74, MKN-45, or MKN-1 cells were seeded in 6 cm dishes (8 × 10^5^ cells per dish) for 24 h in serum-free conditions. Hh signaling mediators were determined and the relative protein expression was compared to SNU-1. (B) Chemical structure of garcinone C. (C) Cytotoxicity of garcinone C in AGS and MKN74 cells was measured after 72 h treatment under serum free conditions. (D) AGS and MKN74 cells were incubated with garcinone C (0.3, 1, 3 μM) for 2 weeks. Colony formation rate was calculated by comparing to the control. The experiment was done in triplicate. All values are presented as means ± SD. Lowercase letters a-d indicate statistically significant differences at p < 0.05 evaluated by one-way ANOVA followed by Duncan’s post hoc test.

### 3.2. Garcinone C induced the G1/G0 phase arrest and apoptosis in gastric cancer cells

Cell proliferation is well known to be regulated by cell cycle and apoptosis^23^. Cell cycle analysis results showed that garcinone C increased the G0/G1 cell population in a dose-dependent manner. In AGS cells, G0/G1 stage increased from 40.7% in the control to 54% with 3 μM garcinone C treatment, while G2/M stage decreased from 37.1% to 23.1%. Similarly, MKN74 cells showed a similar trend (Fig. 2A). In cell cycle, the G1/S and G2/M transitions are key checkpoints controlled by cyclins and cyclin-dependent kinase (CDK) complexes^24^. Cyclin D interacts with CDK4 and CDK6 to drive cell cycle entry, Cyclin E and Cyclin A are able to bind CDK2 to promote G1/S transition^25^. CDK inhibitors like p21, p27, and p57 are main proteins that directly bind to the cyclin-CDK complexes and block their activity^26^. Western blot analysis revealed that 3 μM garcinone C reduced Cyclin D1 levels to 83.0% in AGS and 59.4% in MKN74 cells, while p21 increased more than 2 folds compared to the control. CDK2, the S phase mediator, was not affected by garcinone C treatment in both cell lines (Fig. 2B). Annexin V-PI analysis revealed that garcinone C induced apoptosis in AGS and MKN74 cells (Fig. 2C). The percentage of apoptotic cells increased from 14.6 ± 2.2% to 30.0 ± 0.8% in AGS cells and from 17.4 ± 2.7% to 28.0 ± 0.9% in MKN74 cells following treatment with 3 μM garcinone C. Further analysis of apoptosis-related markers showed that garcinone C increased the levels of pro-apoptotic protein Bax and decreased the anti-apoptotic protein Bcl-2 in both AGS and MKN74 cells (Fig. 2C). It also led to higher levels of cleaved-caspase-3 and cleaved-PARP. Caspase-3, a core protease in the apoptotic process, activating through cleavage and subsequently inactivates poly (ADP-ribose) polymerase (PARP), an endogenous caspase-3 substrate^27^. Cleavage of PARP-1 by caspases is considered to be a hallmark of apoptosis^28^. These results suggest that garcinone C inhibited the proliferation of gastric cancer cells via inducing G0/G1 arrest and apoptosis.

**Figure 2.**
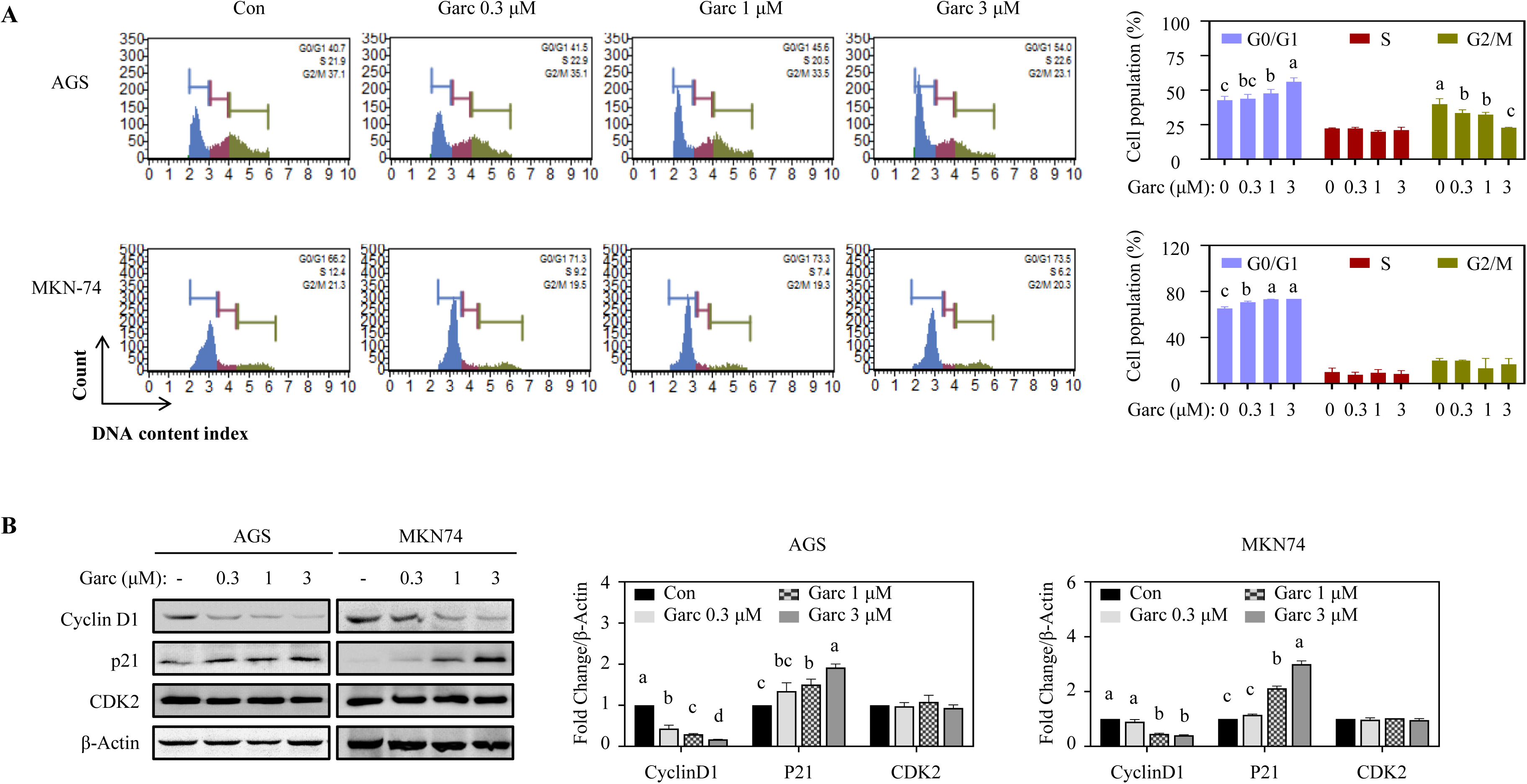
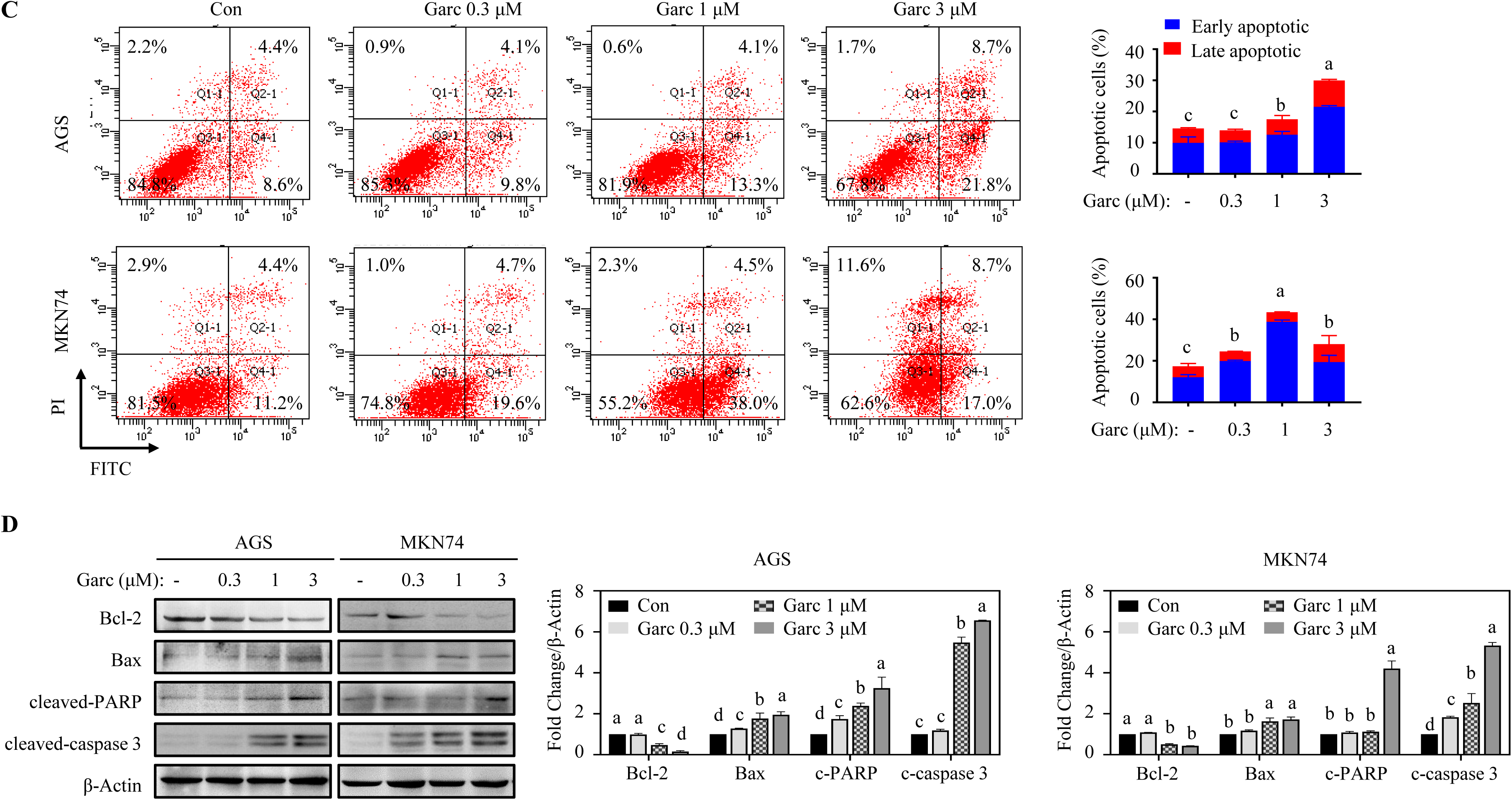
Garcinone C inhibited the proliferation of gastric cancer cells by inducing G1/G0 phase arrest and apoptosis. AGS and MKN74 cells were treated with garcinone C (0.3, 1, or 3 μM) for 24 h in serum-free conditions. Cell cycle analysis was performed using a Muse Cell Analyzer (A) and Annexin V-PI analysis was performed using BD FACSAria II flow cytometer (C). Cell cycle-related (B) and apoptosis-related (D) markers were analyzed by western blot, with relative protein expression levels normalized to β-Actin. The experiment was done in triplicate. All values are presented as means ± SD. Lowercase letters a-d indicate statistically significant differences at p < 0.05 evaluated by one-way ANOVA followed by Duncan’s post hoc test.

### 3.3. Hh signaling inhibitor Gant61 suppressed the proliferation of gastric cancer cells

Previous studies have demonstrated that Hh signaling is involved in the proliferation of gastric cancer *in vitro* and *in vivo*^22^. To explore its role, AGS and MKN74 cells were treated with well-known Hh inhibitors Gant61 (Gli1/2 inhibitor) and vismodegib (Smo inhibitor)^29,30^. Gant61 significantly inhibited cell viability in a dose-dependent manner in both cell lines, while vismodegib had no effect (Fig. 3A). Similarly, colony formation results demonstrated that Gant61 remarkably reduced the colony formation rate to 1.6 ± 1.0% in AGS and 2.8 ± 0.4% in MKN74 cells, while vismodegib had no effect on colony numbers (Fig. 3B). In colon cancer, Gant61 was reported to inhibit the growth of HCT116 cancer cells, whereas vismodegib had no effect^16^. These findings suggest that Gant61 effectively inhibits gastric cancer cell proliferation. In contrast, vismodegib’s lack of effect could be explained by non-canonical, Smo-independent activation of Hh signaling^31^. Smo mutations are the main reason for vismodegib resistance in basal cell carcinoma^32^, and in gastric cancer, galectin-1 promotes metastasis via the Smo-independent Hh/Gli1 pathway^33^. Further supporting this, our results showed that the knockdown of Gli1 or Gli2 reduced cell growth by approximately 30% in both cell lines (Fig. 3C). These results indicate that Smo may be mutated in AGS and MKN74 cells, making Gli1 and Gli2 more effective targets for inhibiting gastric cancer cells.

**Figure 3.**
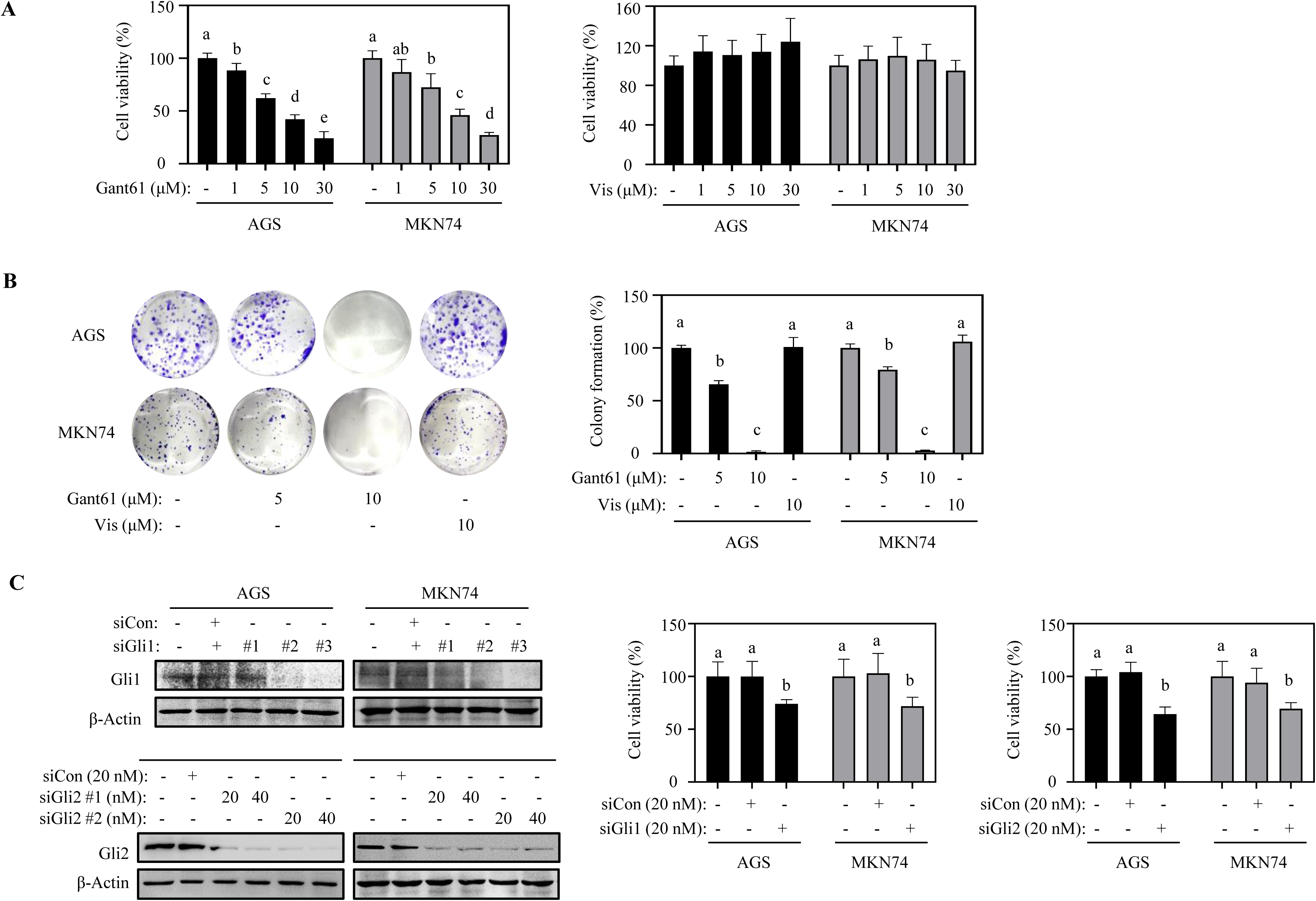
Hh signaling promoted the proliferation of gastric cancer cells. (A) Cytotoxicity of Gant61 and vismodegib in AGS and MKN74 cells was measured after 72 h treatment under serum free conditions. (B) AGS and MKN74 cells were incubated with Gant61 or vismodegib for 2 weeks. Colony formation rate was calculated by comparing to the control. (C) AGS and MKN74 cells transfected with Gli1 or Gli2 siRNA (20 nM) for 24 h in serum-free conditions. Then cells were seeded into 96-well plate (3000 cells/well) and cell viability was checked after 72 h. The experiment was done in triplicate. All values are presented as means ± SD. Lowercase letters a-d indicate statistically significant differences at p < 0.05 evaluated by one-way ANOVA followed by Duncan’s post hoc test.

### 3.4. Inhibiting Hh signaling regulated G0/G1 phase-related markers in gastric cancer cells

Previous studies have reported that proliferation makers such as Cyclin D1, p21, Bax, and Bcl-2 are the direct target of Gli1^34^. Gli1-binding sites have been identified in Cyclin D1 promoter^35^. Morton et al. demonstrated that Shh overexpression enhances pancreatic cancer cell survival from apoptosis by stabilizing Bcl-2 and Bcl-X(L)^36^. Besides, 7 potential Gli binding sites were identified in the Bcl-2 promotor^37^. Hh signaling also modulates Bax phosphorylation at Ser184, changing its role from pro-apoptotic to anti-apoptotin^38^. We evaluated the effect of Hh signaling inhibitors on cyclin D1 and p21. The results showed that only Gant61 significantly down-regulated Cyclin D1 and up-regulated p21 expression levels, while vismodegib had no effect on AGS and MKN74 cells (Fig. 4A). Knockdown of Gli1 with siRNA also found that p21 levels increased more than 2-fold in AGS and 3-fold in MKN74 cells (Fig. 4B). In gastric cancer, knockdown of Gli1 increased p21 levels, disrupted the G1-S transition, and inhibited gastric cancer cell growth^39^. Similarly, Gli1/2 inhibitor Gant61 suppressed neuroblastoma growth by inhibiting Cyclin D1 expression and inducing apoptosis, whereas Smo inhibitor cyclophmine had no effect in neuroblastoma cancer^40^. These studies demonstrate that Gli1-dependent Hh signaling is critical for the proliferation of AGS and MKN74 gastric cancer cells.

**Figure 4.**
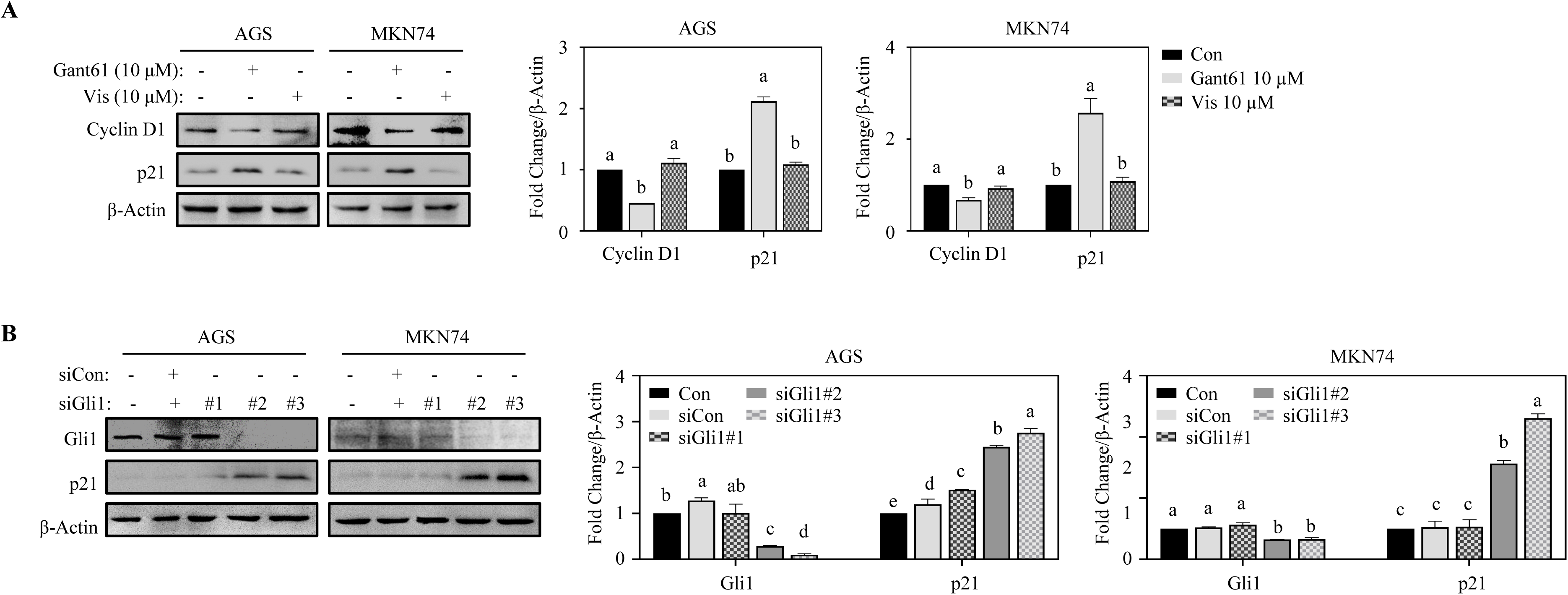
Inhibition of Hh signaling modulated G0/G1 phase-related markers in gastric cancer cells. (A) AGS and MKN74 cells were treated with 10 μM Gant61 or vismodegib for 24 h in serum-free conditions. Cell cycle-related markers were analyzed by western blot, with relative protein expression levels normalized to β-Actin. (B) AGS and MKN74 cells were transfected with Gli1 siRNA (20 nM) for 24 h in serum-free conditions. Cell cycle-related markers were determined by western blot, with relative protein expression levels normalized to β-Actin. The experiment was done in triplicate. All values are presented as means ± SD. Lowercase letters a-d indicate statistically significant differences at p < 0.05 evaluated by one-way ANOVA followed by Duncan’s post hoc test.

### 3.5. Garcinone C inhibited Hh signaling by blocking the nuclear translocation and the protein expression of Gli1/2 in gastric cancer cells

To determine whether garcinone C modulates gastric cancer growth via Hh signaling, we compared its effects with Hh inhibitors. Garcinone C, like Gant61, downregulated Gli1 and Gli2 while upregulating the repressed form of Gli3 in both cell lines (Fig. 5A). However, the Smo inhibitor vismodegib showed no effect. Since the nuclear translocation of Gli1 and Gli2 signifies the complete activation of Hh signaling^18^, garcinone C was found to block the nuclear translocation of Gli1 and Gli2, similar to the effect of Gant61, but not vismodegib in AGS and MKN74 cells (Fig. 5B). It was also confirmed that garcinone C treatment reduced the protein levels of Gli1 and Gli2 in nucleus in both cell lines (Fig. 5C). Previous studies reported that solasodine inhibits Gli1 expression and nuclear translocation through binding to its zinc finger domain, leading to growth inhibition of MCF7 breast cancer stem-like cells^41^. In colorectal cancer, genipin induced apoptosis through Gli1 ubiquitination and suppressed xenograft tumor growth^42^. These findings suggest that garcinone C inhibits gastric cancer cell proliferation through Hh/Gli signaling.

**Figure 5.**
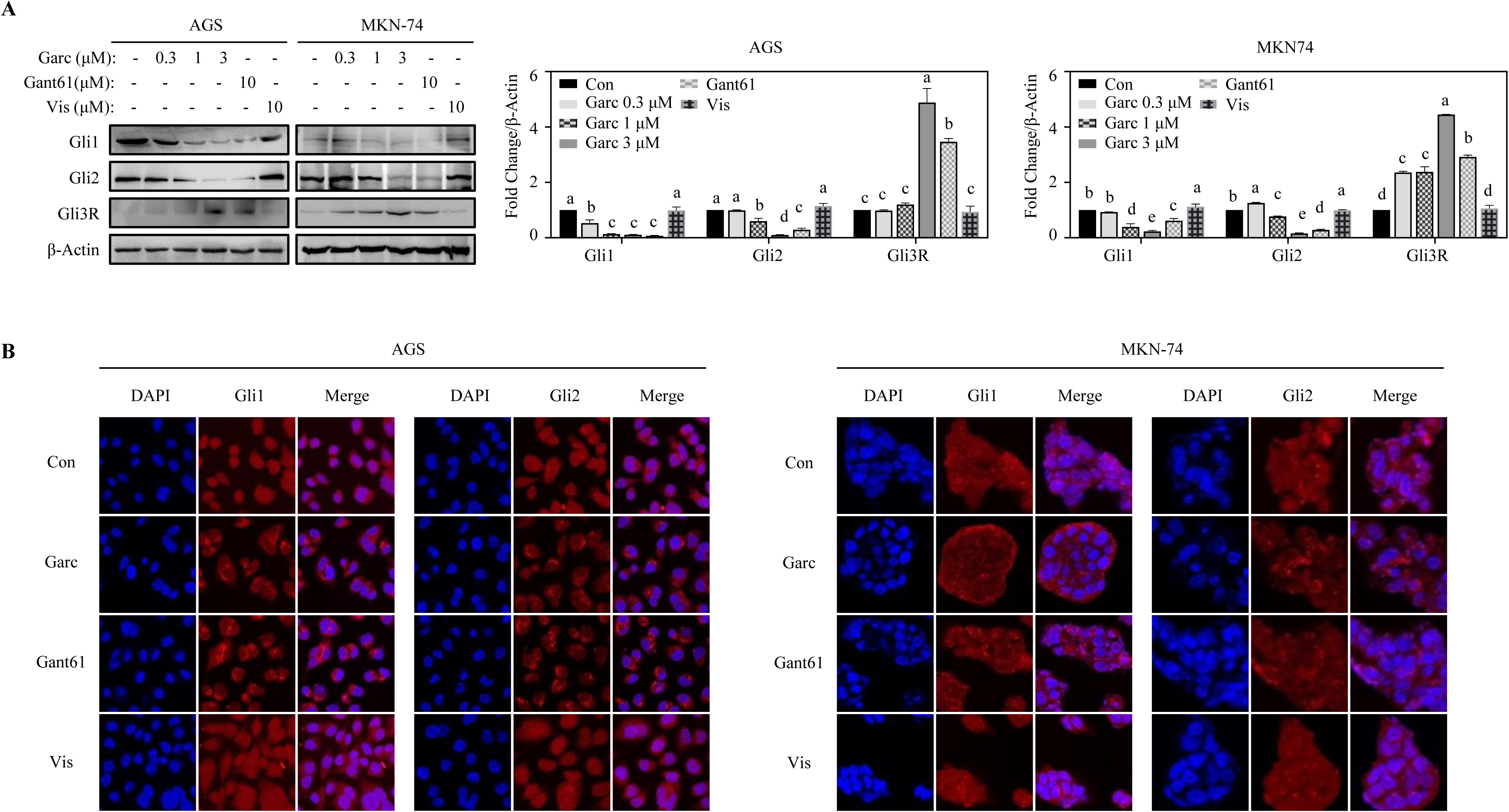
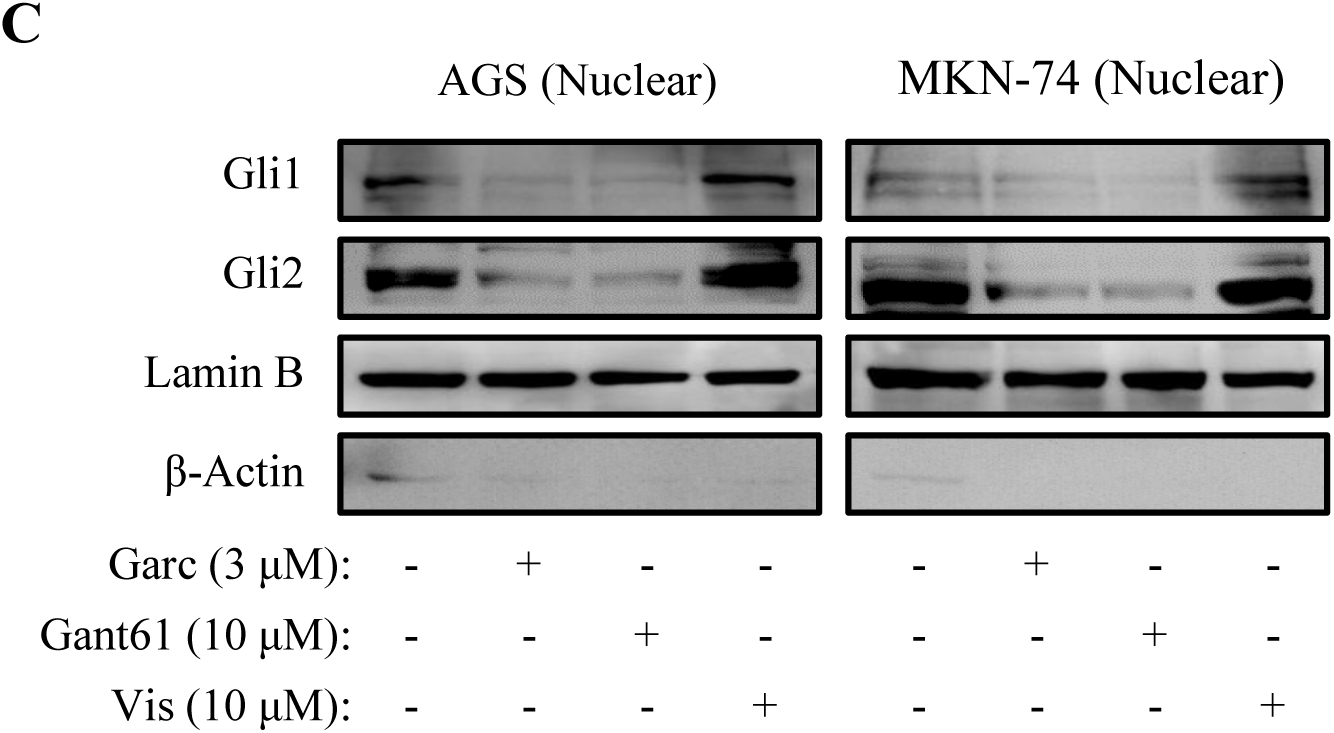
Garcinone C inhibited the proliferation of gastric cancer cells through Hh signaling. (A) AGS and MKN74 cells were treated with garcinone C, Gant61 or vismodegib for 24 h in serum-free conditions. Hh signaling mediators were analyzed by western blot, with relative protein expression levels normalized to β-Actin. (B) AGS and MKN74 cells were treated with garcinone C, Gant61 or vismodegib for 24 h under serum free conditions. Gli1 and Gli2 localization were checked by confocal microscopy (400 × magnification). (C) AGS and MKN74 cells were treated with garcinone C, Gant61 or vismodegib for 24 h in serum-free conditions. The nuclear fractions were collected using the NE-PER extraction kit. Lamin b was used as the nuclear loading control. The experiment was done in triplicate. All values are presented as means ± SD. Lowercase letters a-d indicate statistically significant differences at p < 0.05 evaluated by one-way ANOVA followed by Duncan’s post hoc test.

Garcinone C, a prominent xanthone from mangosteen, has been shown to suppress colon tumorigenesis by blocking Gli1 nuclear translocation and transcriptional activity^16^. Previous studies also demonstrated that 1,3,5,8-tetrahydroxyxanthone suppressed Hh signaling by inhibiting Gli1 expression^43^, further suggesting a role of xanthone in modulating Hh signaling. In line with these findings, our study revealed that garcinone C inhibited the expression and nuclear translocation of Gli1/2, leading to cell cycle arrest, apoptosis, as well as reduced colony formation in AGS and MKN74 gastric cancer cells (Fig. 6). These findings suggest that garcinone C has the potential to inhibit gastric cancer cell proliferation via Hh signaling, highlighting its potential as an anticancer agent. However, for the therapeutic application, it is necessary to validate the effect *in vivo* and the underlying molecular mechanism.

**Figure 6.**
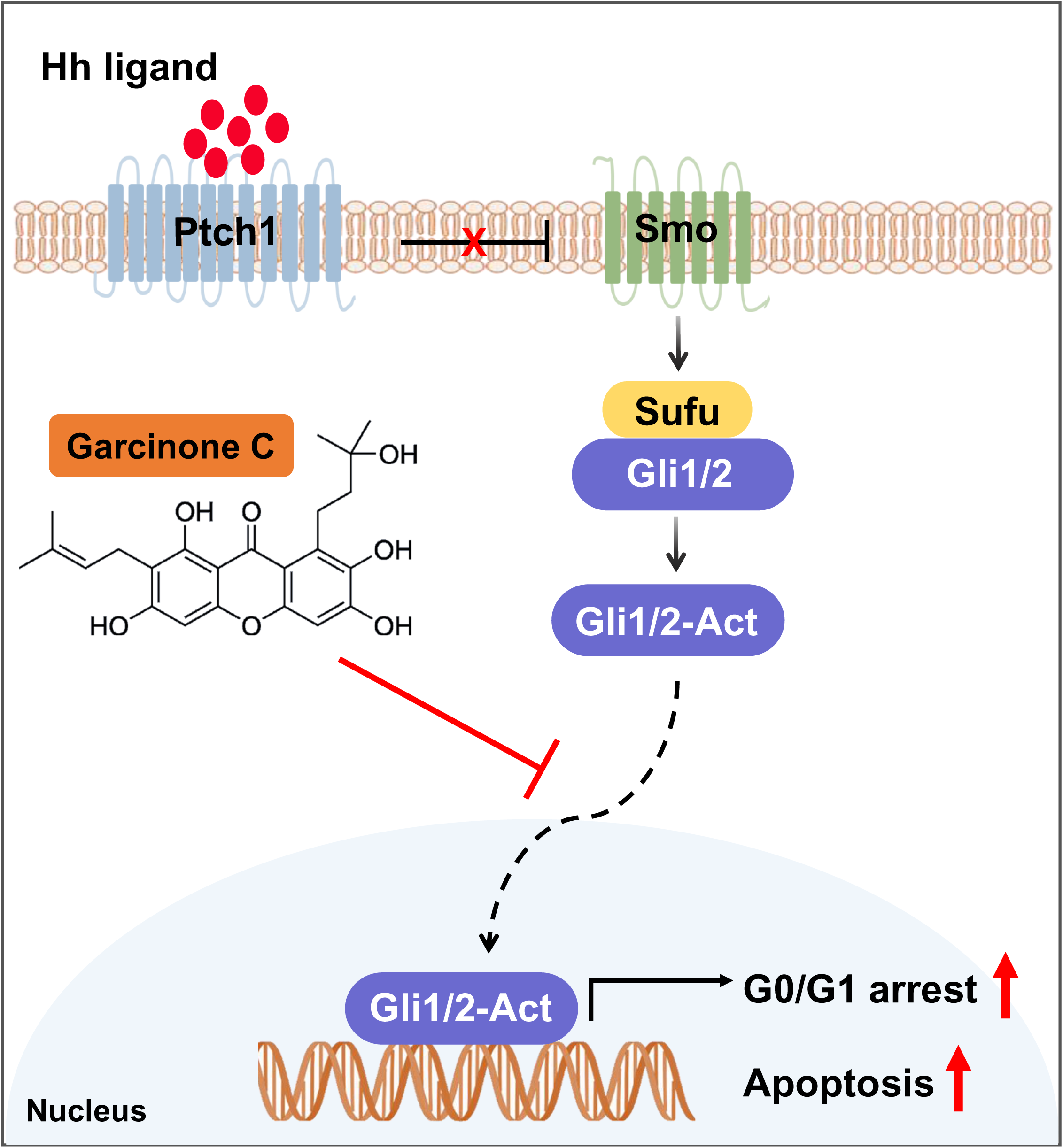
Schematic illustration of the effect of garcinone C on gastric cancer proliferation. In gastric cancer cells, garcinone C suppresses Hh signaling by inhibiting Gli1/2 expression and nuclear translocation, leading to G0/G1 phase arrest and apoptosis, ultimately resulting in reduced cell proliferation.

## 4. Conclusion

In conclusion, we found that garcinone C, a xanthone isolated from *Garcinia mangostana L*, significantly inhibited the proliferation of gastric cancer cells by inducing G0/G1 phase arrest and apoptosis dose-dependently. Inhibiting Gli1 with inhibitor Gant61 or Gli1 siRNA suppressed the growth of AGS and MKN74 cells. Importantly, garcinone C showed similar effects to Gant61 in inhibiting the Hh signaling by suppressing Gli1/2 expression and blocking their nuclear translocation in AGS and MKN74 cells. These findings underscore the potential of garcinone C as a promising therapeutic agent for treating gastric cancer.

## Data availability

The data that support the findings of this study are available on request from the corresponding author.

## Acknowledgments

This research was supported by the National Research Foundation of Korea (NRF-2022R1A2C1092959) and Grants from the Ministry of Food and Drug Safety (21153MFDS605).

## Authors’ contributions

We declare that all authors made fundamental contributions to the manuscript. Conceptualization: Hong Jin Lee, Shuai Qiu; Designed and performed research: Shuai Qiu, Yimeng Zhou, Jin Tae Kim; Data analysis: Shuai Qiu, Yimeng Zhou; Manuscript writing, review and editing: Yimeng Zhou, Shuai Qiu, Jin Tae Kim, Jung Won Kwon, Ga Yeon Lee, Hui Mang Son, Kang Hyuk Lee; Supervision, funding acquisition, and project administration: Hong Jin Lee.

## Reference

1 Guan WL, He Y, Xu RH. Gastric cancer treatment: recent progress and future perspectives. J Hematol Oncol 2023;16:57.

2 Morgan E, Arnold M, Camargo MC, Gini A, Kunzmann AT, Matsuda T, et al. The current and future incidence and mortality of gastric cancer in 185 countries, 2020-40: A population-based modelling study. EClinicalMedicine 2022;47:101404.

3 Janjigian YY, Shitara K, Moehler M, Garrido M, Salman P, Shen L, et al. First-line nivolumab plus chemotherapy versus chemotherapy alone for advanced gastric, gastro-oesophageal junction, and oesophageal adenocarcinoma (CheckMate 649): a randomised, open-label, phase 3 trial. Lancet 2021;398:27–40.

4 Raei N, Behrouz B, Zahri S, Latifi-Navid S. Helicobacter pylori Infection and Dietary Factors Act Synergistically to Promote Gastric Cancer. Asian Pac J Cancer Prev 2016;17:917–21.

5 Molaei F, Forghanifard MM, Fahim Y, Abbaszadegan MR. Molecular Signaling in Tumorigenesis of Gastric Cancer. Iran Biomed J 2018;22:217–30.

6 Zhang J, Fan J, Zeng X, Nie M, Luan J, Wang Y, et al. Hedgehog signaling in gastrointestinal carcinogenesis and the gastrointestinal tumor microenvironment. Acta Pharm Sin B 2021;11:609–20.

7 Wang Z, Lv J, Li X, Lin Q. The flavonoid Astragalin shows anti-tumor activity and inhibits PI3K/AKT signaling in gastric cancer. Chem Biol Drug Des 2021;98:779–86.

8 Zhang X, Zhang C, Ren Z, Zhang F, Xu J, Zhang X, et al. Curcumin Affects Gastric Cancer Cell Migration, Invasion and Cytoskeletal Remodeling Through Gli1-beta-Catenin. Cancer Manag Res 2020;12:3795–806.

9 Wang Y, Wu H, Dong N, Su X, Duan M, Wei Y, et al. Sulforaphane induces S-phase arrest and apoptosis via p53-dependent manner in gastric cancer cells. Sci Rep 2021;11:2504.

10 Akao Y, Nakagawa Y, Nozawa Y. Anti-cancer effects of xanthones from pericarps of mangosteen. Int J Mol Sci 2008;9:355–70.

11 Hung SH, Shen KH, Wu CH, Liu CL, Shih YW. Alpha-mangostin suppresses PC-3 human prostate carcinoma cell metastasis by inhibiting matrix metalloproteinase-2/9 and urokinase-plasminogen expression through the JNK signaling pathway. J Agric Food Chem 2009;57:1291–8.

12 Xu XH, Liu QY, Li T, Liu JL, Chen X, Huang L, et al. Garcinone E induces apoptosis and inhibits migration and invasion in ovarian cancer cells. Sci Rep 2017;7:10718.

13 Li RR, Zeng DY. The effects and mechanism of alpha-mangostin on chemosensitivity of gastric cancer cells. Kaohsiung J Med Sci 2021;37:709–17.

14 Xia Y, Liu X, Zou C, Feng S, Guo H, Yang Y, et al. Garcinone C exerts antitumor activity by modulating the expression of ATR/Stat3/4E-BP1 in nasopharyngeal carcinoma cells. Oncol Rep 2018;39:1485–93.

15 Nauman MC, Tocmo R, Vemu B, Veenstra JP, Johnson JJ. Inhibition of CDK2/CyclinE1 by xanthones from the mangosteen (Garcinia mangostana): a structure-activity relationship study. Nat Prod Res 2021;35:5429–33.

16 Chen J, Qiu S, Kim JT, Cho JS, Moon JH, Zhou Y, et al. Garcinone C suppresses colon tumorigenesis through the Gli1-dependent hedgehog signaling pathway. Phytomedicine 2020;79:153334.

17 Jiang J. Hedgehog signaling mechanism and role in cancer. Semin Cancer Biol 2022;85:107–22.

18 Jiang J, Hui CC. Hedgehog signaling in development and cancer. Dev Cell 2008;15:801–12.

19 Ke B, Wang XN, Liu N, Li B, Wang XJ, Zhang RP, et al. Sonic Hedgehog/Gli1 Signaling Pathway Regulates Cell Migration and Invasion via Induction of Epithelial-to-mesenchymal Transition in Gastric Cancer. J Cancer 2020;11:3932–43.

20 Qiu S, Zhou Y, Kim JT, Bao C, Lee HJ, Chen J. Amentoflavone inhibits tumor necrosis factor-alpha-induced migration and invasion through AKT/mTOR/S6k1/hedgehog signaling in human breast cancer. Food Funct 2021;12:10196–209.

21 Zhou Y, Qiu S, Kim JT, Lee SB, Park HJ, Son MJ, et al. Garcinone C Suppresses Tumorsphere Formation and Invasiveness by Hedgehog/Gli1 Signaling in Colorectal Cancer Stem-like Cells. J Agric Food Chem 2022;70:7941–52.

22 Saze Z, Terashima M, Kogure M, Ohsuka F, Suzuki H, Gotoh M. Activation of the sonic hedgehog pathway and its prognostic impact in patients with gastric cancer. Dig Surg 2012;29:115–23.

23 Alenzi FQ. Links between apoptosis, proliferation and the cell cycle. Br J Biomed Sci 2004;61:99–102.

24 Otto T, Sicinski P. Cell cycle proteins as promising targets in cancer therapy. Nat Rev Cancer 2017;17:93–115.

25 Alao JP. The regulation of cyclin D1 degradation: roles in cancer development and the potential for therapeutic invention. Mol Cancer 2007;6:24.

26 Lai L, Shin GY, Qiu H. The Role of Cell Cycle Regulators in Cell Survival-Dual Functions of Cyclin-Dependent Kinase 20 and p21(Cip1/Waf1). Int J Mol Sci 2020;21.

27 Lyakhovich A, Surralles J. Constitutive activation of caspase-3 and Poly ADP ribose polymerase cleavage in fanconi anemia cells. Mol Cancer Res 2010;8:46–56.

28 Mashimo M, Onishi M, Uno A, Tanimichi A, Nobeyama A, Mori M, et al. The 89-kDa PARP1 cleavage fragment serves as a cytoplasmic PAR carrier to induce AIF-mediated apoptosis. J Biol Chem 2021;296:100046.

29 Agyeman A, Jha BK, Mazumdar T, Houghton JA. Mode and specificity of binding of the small molecule GANT61 to GLI determines inhibition of GLI-DNA binding. Oncotarget 2014;5:4492–503.

30 Robarge KD, Brunton SA, Castanedo GM, Cui Y, Dina MS, Goldsmith R, et al. GDC-0449-a potent inhibitor of the hedgehog pathway. Bioorg Med Chem Lett 2009;19:5576–81.

31 Teperino R, Aberger F, Esterbauer H, Riobo N, Pospisilik JA. Canonical and non-canonical Hedgehog signalling and the control of metabolism. Semin Cell Dev Biol 2014;33:81–92.

32 Atwood SX, Sarin KY, Whitson RJ, Li JR, Kim G, Rezaee M, et al. Smoothened variants explain the majority of drug resistance in basal cell carcinoma. Cancer Cell 2015;27:342–53.

33 Chong Y, Tang D, Gao J, Jiang X, Xu C, Xiong Q, et al. Galectin-1 induces invasion and the epithelial-mesenchymal transition in human gastric cancer cells via non-canonical activation of the hedgehog signaling pathway. Oncotarget 2016;7:83611–26.

34 Zheng X, Zeng W, Gai X, Xu Q, Li C, Liang Z, et al. Role of the Hedgehog pathway in hepatocellular carcinoma (review). Oncol Rep 2013;30:2020–6.

35 Kasper M, Schnidar H, Neill GW, Hanneder M, Klingler S, Blaas L, et al. Selective modulation of Hedgehog/GLI target gene expression by epidermal growth factor signaling in human keratinocytes. Mol Cell Biol 2006;26:6283–98.

36 Morton JP, Mongeau ME, Klimstra DS, Morris JP, Lee YC, Kawaguchi Y, et al. Sonic hedgehog acts at multiple stages during pancreatic tumorigenesis. Proc Natl Acad Sci U S A 2007;104:5103–8.

37 Bigelow RL, Chari NS, Unden AB, Spurgers KB, Lee S, Roop DR, et al. Transcriptional regulation of bcl-2 mediated by the sonic hedgehog signaling pathway through gli-1. J Biol Chem 2004;279:1197–205.

38 Tu YC, Yeh WC, Yu HH, Lee YC, Su BC. Hedgehog Suppresses Paclitaxel Sensitivity by Regulating Akt-Mediated Phosphorylation of Bax in EGFR Wild-Type Non-Small Cell Lung Cancer Cells. Front Pharmacol 2022;13:815308.

39 Ohta M, Tateishi K, Kanai F, Watabe H, Kondo S, Guleng B, et al. p53-Independent negative regulation of p21/cyclin-dependent kinase-interacting protein 1 by the sonic hedgehog-glioma-associated oncogene 1 pathway in gastric carcinoma cells. Cancer Res 2005;65:10822–9.

40 Wickstrom M, Dyberg C, Shimokawa T, Milosevic J, Baryawno N, Fuskevag OM, et al. Targeting the hedgehog signal transduction pathway at the level of GLI inhibits neuroblastoma cell growth in vitro and in vivo. Int J Cancer 2013;132:1516–24.

41 Chen J, Ma D, Zeng C, White LV, Zhang H, Teng Y, et al. Solasodine suppress MCF7 breast cancer stem-like cells via targeting Hedgehog/Gli1. Phytomedicine 2022;107:154448.

42 Kim BR, Jeong YA, Na YJ, Park SH, Jo MJ, Kim JL, et al. Genipin suppresses colorectal cancer cells by inhibiting the Sonic Hedgehog pathway. Oncotarget 2017;8:101952–64.

43 Zhou Y, Kim JT, Qiu S, Lee SB, Park HJ, Soon MJ, et al. 1,3,5,8-Tetrahydroxyxanthone suppressed adipogenesis via activating Hedgehog signaling in 3T3-L1 adipocytes. Food Sci Biotechnol 2022;31:1473–80.

